# Clinically Useful Brain Imaging for Neuropsychiatry: How Can We Get There?

**DOI:** 10.1101/115097

**Authors:** Michael P. Milham, R. Cameron Craddock, Arno Klein

## Abstract

Despite decades of research, visions of transforming neuropsychiatry through the development of brain imaging-based ‘growth charts’ or ‘lab tests’ have remained out of reach. In recent years, there is renewed enthusiasm about the prospect of achieving clinically useful tools capable of aiding the diagnosis and management of neuropsychiatric disorders. The present work explores the basis for this enthusiasm. We assert that there is no single advance that currently has the potential to drive the field of clinical brain imaging forward. Instead, there has been a constellation of advances that, if combined, could lead to the identification of objective brain imaging-based markers of illness. In particular, we focus on advances that are helping to: 1) elucidate the research agenda for biological psychiatry (e.g., neuroscience focus, precision medicine), 2) shift research models for clinical brain imaging (e.g., big data exploration, standardization), 3) break down research silos (e.g., open science, calls for reproducibility and transparency), and 4) improve imaging technologies and methods. While an arduous road remains ahead, these advances are repositioning the brain imaging community for long-term success.

---

As progress is made toward the prevention and treatment of cardiovascular disease and cancers, neuropsychiatric disorders are accounting for an increasing proportion of the unmet disease burden throughout the lifespan (Walker, McGee, and Druss 2015)(Whiteford et al. 2015). Yet our approaches for diagnosing and treating these disorders remain grounded in advances dating back to the late 19^th^ to mid-20^th^ centuries. Over the past two decades, clinical brain imaging has sought to address this gap and has succeeded in revealing biological correlates for a broad array of illnesses (Lagopoulos et al. 2009; Castellanos and Proal 2012; Strakowski, Delbello, and Adler 2005; Etkin and Wager 2007; Savitz, Rauch, and Drevets 2013;Rane et al. 2015). However, the inability to find clear relationships between abnormal brain function and traditional diagnostic classifications of neuropsychiatric illness has hindered the pace of progress. In particular, it has resulted in an inadequate biological understanding of these illnesses and difficulty for imaging to: 1) identify meaningful biological indices of clinical diagnosis, prognosis or risk (i.e., biomarkers), and 2) facilitate the development of novel targets for therapeutic interventions (Kapur, Phillips, and Insel 2012). In short, despite decades of effort, visions of transforming clinical practice for neuropsychiatric illness through developing objective imaging-based ‘growth charts’ or ‘lab tests’ for the brain have remained out of reach.

In recent years, efforts focused on mapping the human connectome (the wiring diagram of the human brain) (Craddock et al. 2013)(D. C.Van Essen and Ugurbil 2012) and its functional interactions (Kelly et al. 2012) have sparked renewed enthusiasm about the possibility of realizing clinically useful biomarkers for neuropsychiatric illness. We consider a biomarker to have clinical utility if it not only correlates with the presence of an illness, but can help to inform diagnosis, treatment selection/evaluation, assessments of risk, and prognosis (Rubinov and Bullmore 2013)(Fornito and Bullmore 2015)(Fox and Greicius 2010). This renewed enthusiasm is not without its detractors, who ask what is different from the past, what is the ‘game changer’? Here, we argue that there is no single game changer. Instead, we review a constellation of advances that together improve the chances of delivering biological markers of illness for neuropsychiatric disorders.

## Elucidation of the research agenda for biological psychiatry

The first key question for any pursuit focused on developing clinically useful measures from brain imaging is: *what are we trying to accomplish?* The vast majority of clinical brain imaging research has focused on extreme comparisons between groups of individuals with a clean Diagnostic and Statistical Manual of Mental Disorders (DSM–5) diagnosis (such as major depressive disorder with no comorbidities) and "pure” controls (Kapur, Phillips, and Insel 2012). Such studies do not reflect the "real world” challenges for the assessment of neuropsychiatric disorders. Nearly half of all individuals affected by mental illness meet criteria for two or more mental health disorders (Bijl, Ravelli, and van Zessen 1998), and DSM classifications for these disorders likely include a heterogenous mix of biological profiles. The real power of brain imaging will be in differentiating seemingly overlapping clinical disorders, as well as subtypes of the same disorder, rather than simply differentiating typical individuals from those affected by one or more disorders. This ability to differentiate could also help to improve the assignment, targeting, and monitoring of treatment interventions, and provides a more pragmatic scenario for translating brain imaging research to the clinic.

### A shift away from disorder-focused definitions of psychiatry

The biological psychiatry community has come to accept the limitations of efforts to map current psychiatric nosologies onto the brain (T. Insel et al. 2010; Kapur, Phillips, and Insel 2012). Although a logical starting point for the field, efforts to map classifications in current diagnostic manuals (e.g., DSM 5) to the brain, or to other biological modalities (e.g., genes) have had mixed success. The field has yet to find reproducible and generalizable one-to-one mappings between diagnoses and patterns of brain abnormality. This is not surprising, as these disorders were defined based on the detection of clusters of signs and symptoms of psychiatric illness, not based on biology.

While frustrating, this lack of specificity has given rise to a new agenda for the biological psychiatry community - namely, the development of a neuroscience-based diagnostic classification system for mental health and learning disorders. The NIMH Research Domain Criteria project (RDoC) has taken a lead role in pushing a neuroscience-focused agenda for neuropsychiatry (Cuthbert and Insel 2013). The calls for an increased neuroscience focus are not necessarily intended to be a complete rejection of classical nosologies, but rather, a challenge to rethink the existing diagnostic boundaries. Within a given domain of illness such as ADHD, researchers are being asked to find biologically-based subtypes that can help explain variation among affected individuals. These subtypes would have the potential to be more informative than the rudimentary subdivisions defined by clinical observation, such as inattentive versus hyperactive-impulsive versus combined for ADHD (Gates et al. 2014).

Researchers are also being challenged to use neuroscience to rethink existing boundaries for domains of illness that extend across multiple diagnostic categories, such as depression (e.g., bipolar, unipolar, atypical, and chronic) (T. R. Insel and Cuthbert 2015). For example, a recent review by (McTeague, Goodkind, and Etkin 2016) highlighted the presence of broad perturbations in behavioral indices of cognitive control for a range of disorders (mood, anxiety, psychotic and substance use disorders); they argued that only the severity of the perturbations observed appears to depend on the specific disorder being examined. Consistent with these behavioral findings, imaging studies have reported abnormalities within the frontoparietal multiple demands system (Duncan 2010) across the range of disorders, particularly in dorsal anterior cingulate and bilateral insular cortices. Such observations motivate further research that looks across diagnostic labels in clinically diverse samples for commonalities in impairments and in biology (see (Van Dam et al. 2016) for another recent example); this research in turn has the potential to guide development of transdiagnostic treatments for impairments of interest.

Researchers are also being challenged to consider comorbid illnesses, which too often are overlooked in applications of the current diagnostic nosology in brain imaging research, despite undeniably high rates. For example, a recent US-based study found the following comorbidities for adults with ADHD (Willcutt 2012): social phobia: 29.3%, specific phobia: 22.7%, bipolar disorder: 19.4%, major depressive disorder: 18.6%, intermittent explosive disorder: 19.6%, and substance use disorder: 15.2%. The integration of neuroscientific data will allow us to determine when seemingly disparate symptoms are actually linked to a common biological origin, and when seemingly similar phenomena have different biological origins.

### The emerging concept of precision medicine

Over the past five years, the concept of precision medicine has emerged as a guiding approach for the future of healthcare research and practice (Collins and Varmus 2015)(Mirnezami, Nicholson, and Darzi 2012). The medical community has traditionally relied on the use of a one-size-fits-all approach to the prevention and treatment of an identified disease, where the same treatment is applied to all individuals with a given diagnosis. Unfortunately, such strategies fail to take into account inter-individual variability, resulting in a trial and error approach to figuring out who will respond well to a given intervention and who will not, as well as who will have an adverse reaction and who will not. Precision medicine attempts to minimize the trial and error aspect of interventions by stratifying individuals based on one or more factors that are predictive of response. Candidate predictors that could potentially guide stratification include biological factors such as genes, immunological profiles, and imaging findings, as well as lifestyle and environmental exposures. It is worth noting that precision medicine differs from the concept of personalized medicine in that it is not calling for us to engineer treatments for the individual, but rather to select from existing treatments the one that is most likely to work for an individual (Sipka 2016). Arguably, this is a more attainable goal. A demonstration of the potential role for brain imaging in precision medicine comes from a recent study that was able to: a) identify four biological subtypes of depression in a sample of 711 individuals using resting-state functional magnetic resonance imaging (fMRI) measures of brain connectivity, and b) use subtype information to predict responsiveness to transcranial magnetic stimulation therapy (Drysdale et al. 2017).

By emphasizing stratification, proponents of precision medicine have fueled ambitions to make brain imaging more clinically useful. Psychiatric researchers have long sought to develop clinical tools that could be helpful in the diagnostic process for complex cases, much the way a strep test can be used to disambiguate the causes of a sore throat in primary care settings and guide treatment. Stratification heralds a broader research agenda that focuses on the assessment of prognosis (e.g., likelihood of disease progression or treatment response) and risk (e.g., likelihood of developing a disease or comorbid disease). The development of objective measures of risk and prognosis are essential for efforts focused on early intervention or prevention.

While the prospect of bringing the precision medicine model to psychiatry is a source of excitement (Kapur, Phillips, and Insel 2012; T. R. Insel and Cuthbert 2015;Hahn, Nierenberg, and Whitfield-Gabrieli 2016), a number of logistical and infrastructural challenges exist for its implementation (Joyner and Paneth 2017). The biggest (and most costly) challenge is obtaining longitudinal data from individuals to identify predictors of clinically relevant outcomes, especially in dedicated research studies. An attractive solution is to leverage the longitudinal clinical data obtained in clinical settings (past and future) for the purposes of research; however, the implementation of such strategies is hindered by privacy concerns, poor data quality and limitations in interoperability among differing electronic health record systems. Efforts such as the NIH-funded Informatics for Integrating Biology and the Bedside (i2b2; https://www.i2b2.org/)(Weber et al. 2009) are working to bridge these gaps and facilitate discovery efforts (see (Lingren T 2017) and (Doshi-Velez F 2017) for examples).

## Shifting research models for clinical brain imaging

### Bringing the individual into focus

There is a key confusion in the literature between brain-based differences that are informative to scientists and those that are useful to clinicians. Scientific tradition has long established the pursuit of statistical significance as the goal of identifying differences among individuals, with the threshold set at a minimum standard of p < 0.05 (following appropriate statistical correction for multiple comparisons). Consistent with this tradition, myriad clinical brain imaging studies have worked to demonstrate statistically significant brain differences between affected individuals and comparisons. These differences have proven to be highly useful in the generation and testing of scientific models of psychiatric illness, and are well-justified for such pursuits. However, as highlighted by several recent works (Kapur, Phillips, and Insel 2012)(Castellanos et al. 2013), population-level differences in brain or behavior indices have little direct clinical applicability, as they are not sufficiently informative about the individual standing in front of a clinician. A far more challenging set of standards is required to derive tools that can be used for clinical purposes. As highlighted in a recent review (Castellanos et al. 2013), effect sizes required for a clinically useful biomarker can range between a Cohen's d of 1.5 and 3.0 depending on the nature of the application (e.g., screening, diagnosis). Such effect sizes are rare in the clinical brain imaging literature (Müller et al. 2017; Thompson et al. 2017; Schmaal et al. 2016), though they represent the true goal for which studies focused on clinical applications should strive - not the more traditional pursuit of p < 0.05 significance. Consistent with the requirements of any laboratory test, sufficient accuracy/validity, reliability/precision, sensitivity and specificity must also be established for brain-based biomarkers before any conversation of clinical application should begin.

In recent years, the increasing popularity of machine learning in the imaging community (Pereira, Mitchell, and Botvinick 2009; Lemm et al. 2011; Davatzikos et al. 2005) has brought with it shifts in the design of experiments from testing for clinical differences among populations to individual-level prediction of clinical variables, such as diagnostic status. For those focused on the development of clinical tools, this shift is necessary, as it brings the target closer to direct clinical applications. However, as highlighted in greater detail elsewhere (Castellanos et al. 2013), care needs to be taken in the handling of findings from prediction-based studies, as a number of factors can limit their readiness for clinical application. Such factors include the level of rigor involved in cross-validation approaches (e.g., leave-one-out, k-fold, split-half, and independent-sample) (Varoquaux and Craddock 2013) and how well a study sample reflects the real-world characteristics of a diseased population.

### The big data research model

Following decades of innovations in technology and methods, the neuroscience of the 21st century is increasingly being defined by large-scale efforts to understand complex brain systems using massive datasets (Sejnowski, Churchland, and Movshon 2014; Kandel et al. 2013; Poldrack and Gorgolewski 2014) (Craddock, Tungaraza, and Milham 2015). This approach to scientific investigation, referred to as the "big data" research model, has already yielded unprecedented results in diverse domains such as genetics, physics, astronomy and medicine. The brain imaging community – which historically has been criticized for its reliance on underpowered samples, approximate replications and a research silo culture – has emerged as an entry point for the big data model in neuroscience.

The implications of the big data model are multifold for the future of clinical brain imaging. First, in contrast to the more traditional scientific model ingrained in the imaging community, questions, not hypotheses, are the guiding light for investigations. The exploratory focus of the big data model is not intended to minimize the value of hypothesis-driven research. Instead, it is intended to be a complementary approach, capable of leading to the formation of novel hypotheses through the detection of previously unknown associations. The big data model also enables researchers to look for multivariate relationships which may not be readily accessible with more traditional models (Varoquaux and Craddock 2013). The application of unsupervised learning methods (e.g., cluster analysis) to neuroscientific data can be used to identify previously unappreciated groupings of individuals within a given domain of illness (e.g., novel subtypes for ADHD or depression), or to build links across behaviorally defined boundaries that do not reflect known underlying neurobiology (Gates et al. 2014; Miranda-Dominguez et al. 2014). Supervised learning techniques can be used to develop predictive models for precision medicine (Craddock et al. 2009; Castellanos et al. 2013).

It is important to note that from a clinical application perspective, the value of large-scale datasets is not their ability to increase statistical power. Our ability to detect small to moderate effects will undoubtedly increase with sample size; while these can be valuable for appreciating scientifically interesting population-level associations, as previously discussed, clinical utility hinges instead on the detection of large effects that can be used to meaningfully generate predictions about an individual. As such, the value of big data samples for clinical imaging lies in their ability to represent the broader range of clinical and biological heterogeneity in populations of interest, which is essential for creating more generalizable and informative research tools. Additionally, large sample sizes enable more rigorous cross-validation approaches, such as split-half cross-validation(Strother et al. 2002).

### Rethinking phenotyping

Arguably, the most valuable aspect of brain imaging data is actually the associated phenotypic data; a given anatomical, diffusion or resting-state fMRI dataset has little value if we don’t know anything about the individual from which the images were acquired or the context within which they were acquired. Unfortunately, many researchers are only beginning to realize the importance of careful phenotyping. While the community has focused on the reliability of imaging measures for years, many cognitive and behavioral measures are brought directly into imaging studies without sufficiently establishing their reliability. Consistency in demographic, cognitive, and behavioral measures used across studies is also a challenging issue resulting from the lack of standards; for example, a multitude of questionnaires exist, each providing slightly different information. In response to these challenges, there is a growing focus on the development of standardized data elements that can be included in all studies, known as ‘common data elements’ ("General Standards - NINDS Common Data Elements”)(Simmons and Quinn 2014)(Thurmond et al. 2010), as well as standardized (semi-structured and structured) clinical interviews with defined mechanisms for establishing inter-rater reliability (e.g., ADOS (Lord et al. 2008) for autism).

A common alternative to the use of DSM labels for the characterization of phenotypic variation is the use of dimensional psychiatric assessments (e.g., (Achenbach 2016)). Typically reliant on questionnaires, the tendency of such approaches is to probe for the presence or absence of symptoms. However, this practice frequently results in the creation of truncated distributions, with little variance among those individuals who are without symptoms. Recent efforts are attempting to draw attention to the value of designing instruments that can index both strengths and weaknesses related to particular domains of function. For example, the Strengths and Weaknesses of ADHD Symptoms and Normal Behavior Rating Scales (SWAN) is a rating scale that was explicitly designed to index the broader range of attentional function across individuals, yielding near-normal distributions (Hay et al. 2007; Arnett et al. 2013; Lakes, Swanson, and Riggs 2012); such distributions are optimal for application in population studies. The Extended Strengths and Weaknesses Assessment of Normal Behavior (E-SWAN; http://eswan.org) is currently under development to extend the design concept from the SWAN to a broader range of psychiatric domains.

Researchers are also beginning to turn their attention to mobile technologies such as smartphone applications (Bot et al. 2016) and wearable devices to collect cognitive, behavioral, mood, and physiological data using active and passive data collection methods. Active methods include explicit self reports, from occasional and detailed survey instruments to more frequent and brief interactions, sometimes referred to as "ecological momentary assessments,” as well as the performance of tasks probing various domains of function (e.g., memory). Passive methods usually involve sensors worn on some part of the body that can unobtrusively collect high temporal frequency data (e.g., accelerometry actigraphs, photoplethysmography, electrodermal activity, electromyography, etc.). Passive methods can collect objective, real-world data about participants to track cognitive, behavioral, mood, and physiological states, detect medication response, etc. (Chaibub Neto E 2017), in ways that could never be achieved before (Glenn and Monteith 2014; Maetzler et al. 2013)(Adams et al. 2016). Collecting and analyzing these different sources of data help to foster a high-dimensional view of health.

## From research silos to a global scientific community

As biological psychiatry research adopts an agenda with a greater focus on neuroscience and precision medicine, and an emphasis on research models that collect and analyze larger and more varied datasets, new challenges arise. In particular, increased demands for sophisticated informatics and analysis resources far surpass the capabilities of any single investigative team, consortium or scientific discipline (Neuro Cloud Consortium 2016). The field can overcome such seemingly insurmountable obstacles in the realization of clinically useful biomarkers by shifting away from the status quo, where discovery is hampered by the limited resources and capabilities of competing research entities, to a collaborative model that leverages the combined strengths of the research field as a whole. Widescale, effective collaboration demands significant sharing, which is in line with the tenets of open science - a commitment to providing unrestricted access to all ideas, data, tools, and other artifacts generated in the scientific process (Carvalho 2015). In recognition of the benefits of this shift, funding institutions and research organizations are turning to, and in some cases mandating, open science as a means of addressing the massive requirements of this enterprise.

### Scaling up data resources

The natural prerequisite to successful implementation of the big data model is the accrual of massive-scale, heterogeneous and deeply phenotyped datasets. Few labs can hope to generate more than a hundred high quality, well-phenotyped brain image datasets in a year, which sets an unreasonably lengthy timeline for the big data agenda. However, the open science model can make very large datasets available much more quickly, and in many cases more cost effectively, by pooling data collected from many different laboratories. Data sharing efforts such as the 1000 Functional Connectomes Project and its International Neuroimaging Data-sharing Initiative (Mennes et al. 2013) (FCP/INDI), ADNI (Mueller et al. 2005/7) and OpenFMRI (Poldrack et al. 2013) are quickly increasing the scale of analyses that can be performed by the brain imaging community. FCP/INDI projects focused on specific disorders, such as the Autism Brain Imaging Data Exchange (ABIDE) and ADHD-200 are testaments to the type of resources that can be aggregated when researchers in the same topic area collaborate rather than compete.

It is worth noting that the large datasets generated by successful data sharing initiatives require considerable image processing, making them potentially unusable by researchers with limited access to computational expertise and resources. One solution to this challenge is the creation of collaborative efforts such as the Preprocessed Connectomes Project (http://preprocessed-connectomes-proiect.org/), in which researchers who have these computational resources prospectively collaborate with the entire research field by preprocessing the data and sharing the results (Bellec et al. 2016). Data processing steps that require manual interventions can also be handled much more quickly and efficiently if performed collaboratively by a large number of researchers using crowdsourcing platforms such as Brain Box (http://www.brainbox.co.uk/) or through online games such as Eyewire (http://eyewire.org/).

### Directing the challenge to the broader scientific community

Beyond bolstering the efforts of existing community members, open science initiatives remove barriers to entry for members of the broader scientific community. Large-scale brain imaging data are rife with complexities that push the boundaries of what is possible with conventional inferential statistics. Given the need for advanced analysis methods and infrastructures that are outside of the areas of expertise for most clinician scientists, it essential to recruit computer scientists, engineers, mathematicians, and statisticians to mental health research. The ADHD-200 (Milham et al. 2012) and ABIDE (A. Di Martino et al. 2014) initiatives demonstrated that making data available from a population with a given condition draws focus to that condition and engages individuals from a range of disciplines. Contests such as the ADHD-200 Global Competition or the Alzheimer's Disease Big Data DREAM Challenge (https://www.synapse.Org/Synapse:syn2290704) can help to accelerate the recruitment of investigators from other disciplines (Allen et al. 2016)(Milham 2012). The integration of researchers from other domains requires education and training resources, however. Those from non-imaging, computational disciplines will need to be educated on the fundamentals of brain imaging data, while clinical researchers will need to improve their computational and analytical skills to promote the cohesiveness and quality of their research. This is being addressed by educational workshops such as Brainhack (Craddock et al. 2016) and Neurohackweek (“Neurohackweek” 2016), where interdisciplinary researchers come together to learn from one another as they work to address open problems in neuroscience.

### Increasing the focus on reproducible research and transparency

Current estimates suggest that as many as 50% of scientific findings cannot be reproduced (Ioannidis 2005). Excepting rare cases of blatant fraud, the lack of reproducibility in research can be attributed to a variety of factors, ranging from underpowered sample sizes, differences in experimental methods, and heterogeneous samples, to liberal statistical thresholding, insufficient descriptions of study procedures and errors in in-house software (Müller et al. 2017; Eklund, Nichols, and Knutsson 2016; Woo, Krishnan, and Wager 2014). Open science is increasingly heralded as a solution to these challenges. Concepts such as open data sharing, open source software, and open lab notebooks are promising to revolutionize research practices and dramatically augment reproducibility. From a practical view, such levels of transparency are likely to increase the standard of data quality and documentation; researchers will know that others will scrutinize their work, and have the opportunity to improve upon the original work through feedback and experimentation. Even the publication process is beginning to be transformed by open science, as preprint archives such as arXiv, bioRxiv, and Authorea are encouraging authors to upload their manuscripts for viewing soon after submission, rather than at the end of a review process -- a practice which is dramatically speeding up the communication of findings. For example, the manuscript by Eklund et al., 2016(Eklund, Nichols, and Knutsson 2016) that raised concerns about the accuracy of commonly used statistical corrections for fMRI was openly available as a preprint one year prior to publication. Venues such as Academic Karma, Frontiers, Gigascience and F1000 are pushing the idea of open review, where reviewers no longer blind themselves, removing a commonly cited source of bias in the publication process.

Although promising, the implementation of open science is not without its challenges. In particular, the field is still working through issues related to protection of privacy for high-dimensional datasets to ensure the safety of research participants. Mechanisms are still being worked out to promote academic accreditation in an open science community (e.g., data papers (Gorgolewski, Margulies, and Milham 2013)), as current tenure promotion practices tend to engender feelings of competition rather than promote sharing. The recent increase in attention to open science by funding agencies such as the NIH and NSF will likely prove to be the greatest catalyst to motivate change at all levels.

## Breaking down barriers by advancing methods and technologies

### The maturation of pediatric brain imaging

The majority of neuropsychiatric illnesses have onsets in the first two decades of life (Kessler et al. 2005), when brain structure and function are actively developing and maturing. As such, the broader range of neuropsychiatric disorders is increasingly being conceptualized as neurodevelopmental disorders (Adriana Di Martino et al. 2014), with origins possibly tracing back as early as the fetal and perinatal periods (O’Donnell and Meaney 2016)(Schlotz and Phillips 2009). It follows that early detection and interventions, as well as prevention-focused efforts, could necessitate early-life and childhood imaging (Koyama et al. 2016). As discussed in greater detail elsewhere (Adriana Di Martino et al. 2014)(van den Heuvel and Thomason 2016), advances in task-independent brain imaging methods (e.g., resting-state fMRI, morphometry MRI, diffusion MRI) are rapidly expanding the window of investigations focused on human brain function to early portions of the lifespan, including toddler, infant, neonatal and fetal imaging. This expansion is invigorating the developmental imaging community and increasing enthusiasm about the potential to develop tools capable of guiding early intervention efforts for disorders such as autism and schizophrenia.

### Increasing harmonization of methods

Previously described efforts such as the FCP/INDI have provided examples of the potential to quickly amass large-scale datasets through the aggregation of data independently collected at different research facilities. However, the lack of coordination in the acquisition of imaging (and phenotypic) data can introduce site-related variation in the data that can limit our ability to detect underlying biological signals. To address this challenge, recent efforts have demonstrated the potential value of accounting for unintended site-related variation through statistical corrections (Keshavan et al. 2016), or explicitly optimizing image analysis pipelines and techniques to maximize the reproducibility of findings across different imaging sites (Abraham et al., n.d.)(Eloyan, Crainiceanu, and Caffo 2013)(Yan, Craddock, Zuo, et al. 2013)(Fortin et al. 2016) (Abraham et al. in press).

Looking forward, prospective large-scale data collection consortia such as ADNI (Mueller et al. 2005/7), the Brain Genomics Superstruct (Holmes et al. 2015), the upcoming NIH Human Connectome Lifespan Project (David C. Van Essen 2011), PING (Jernigan et al. 2016), and the NIH ABCD Study (“ABCD Study” 2016) have worked to address the challenges of imaging site-related variation through harmonizing their acquisition protocols, as well as the use of phantoms for calibrating data between sites. Among these efforts, ADNI, PING, and the NIH ABCD Study have taken on the particular challenge of harmonizing protocols across different MRI scanner platforms. The success of these efforts is leading to calls in the brain imaging community for increased standardization of data acquisition protocols across sites in general. While the provision of basic guidelines for acquisition can improve the overall quality and comparability of data, the benefits need to be balanced against the need for continued efforts to innovate data acquisition techniques; likely such decisions should be made on a case-by-case basis. At a minimum, encouraging and supporting the routine acquisition of data from standardized phantoms can facilitate efforts to calibrate data across scanner sites (Gunter JL 2017)(Gouttard et al. 2008)(Jovicich J 2017; Keenan et al. 2016)(DeDora et al. 2016).

### Improving MRI scanner capabilities

Modern advances in scanning technology are increasing the quality, resolution, and efficiency of data collection. Advances in head coil technologies have improved SNR and have enabled higher factors of parallel imaging, which combine to increase the resolution and speed for structural imaging acquisition. Newly developed simultaneous multislice (SMS) imaging sequences have increased the temporal and spatial resolutions achievable for fMRI (Feinberg and Setsompop 2013). SMS combined with stronger magnetic field gradients are vastly improving the quality of diffusion MRI data that can be acquired during routine clinical evaluations. These recent significant gains in MRI technology serve as a valuable reminder that MRI is still a young technology and there are still opportunities for improvement.

In particular, continued innovation in MRI scanning equipment and sequences will be essential to fully realize the potential of brain imaging biomarkers and revolutionize clinical practice. Head motion is the nemesis of all brain imaging applications, but is particularly problematic when relying on quantitative measurements from the data (Van Dijk, Sabuncu, and Buckner 2011; Friston et al. 1996; Reuter et al. 2015). Methods for prospectively correcting head motion in structural MRI are showing promise (Tisdall et al. 2012), but their efficacy in clinical settings has yet to be proven. For fMRI, while a motion-robust scanning sequence has yet to emerge, a variety of post-hoc correction techniques have been proposed and evaluated (Power et al. 2012; Yan, Craddock, He, et al. 2013; Pruim et al. 2015; Friston et al. 1996). The challenges of head motion for diffusion MRI are similar to fMRI, and as such are becoming an increasing area of focus for future work (Yendiki et al. 2014).

### Improved methods for large-scale image processing and analysis

With every advance in scanning technologies and increase of scale in data analysis comes the need for new and improved data processing and analysis tools. The growing availability of large-scale datasets is bringing about a change in the design of brain imaging tools, which up to now have focused primarily on the needs of smaller scale datasets (Craddock, Tungaraza, and Milham 2015). Shortcomings of parametric models of brain imaging data have also highlighted the need to switch to more computationally intensive Monte Carlo methods for assessing statistical significance (Eklund, Nichols, and Knutsson 2016; Eklund et al. 2012). To make this computation possible, a new emphasis is being placed on optimizing image processing to minimize its memory footprint and computational demands. Tools that implement and integrate with graphical processing units, cloud computing, and other computational infrastructure are emerging as potential solutions to the challenge of providing relatively low-cost computational resource options to users who do not necessarily have such expertise (Eklund, Andersson, and Knutsson 2012). Making this work at a large scale will require a shift from relying exclusively on local collaborations to designing software that takes care of these details for the user.

## Concluding remarks

We assert that there is no single advance that currently has the potential to drive the field of clinical neuroscience forward. Instead, there has been a constellation of advances that, if combined, could bring about the insights, momentum, and direction to discover biomarkers of neuropsychiatric illness. While our focus was on advances that promise to transform multimodal imaging studies, there is a broader landscape of change and innovation across the range of disciplines focused on biomarker discovery (e.g., genetics, immunology, electron microscopy). Over time, the integration of these varied fronts may prove to bring about the most pivotal changes yet for clinical neuroscience.

## Acknowledgements

This work was supported by gifts to the Child Mind Institute from Phyllis Green, Randolph Cowen, and Joseph P. Healey.

